# Mitochondrial DNA SNPs associated with Schizophrenia exhibit Highly Variable Inter-allelic Haplogroup Affiliation and Nuclear Genogeographic Affinity: Bi-Genomic Linkage Disequilibrium raises Major Concerns for Link to Disease

**DOI:** 10.1101/149070

**Authors:** Christian M Hagen, Vanessa F Gonçalves, Paula L Hedley, Jonas Bybjerg-Grauholm, Marie Bækvad-Hansen, Christine S Hansen, Jørgen K Kanters, Jimmi Nielsen, Ole Mors, Alfonso B Demur, Thomas D Als, Merete Nordentoft, Anders Børglum, Preben Bo Mortensen, James Kennedy, Thomas M Werge, David M Hougaard, Michael Christiansen

**Affiliations:** Department for Congenital Disorders, Statens Serum Institut, Copenhagen, Denmark; Centre for Addiction and Mental Health, University of Toronto, Toronto, Canada; Department of Biomedical Sciences, University of Copenhagen, Copenhagen, Denmark; Aalborg Psychiatric Hospital. Aalborg University Hospital, Aalborg, Denmark; Department of Clinical Medicine, Aarhus University, Århus, Denmark; Mental Health Centre, Sct Hans, Capital Region of Denmark, Denmark; Institute of Medical Genetics, Aarhus University, Århus, Denmark; Mental Health Centre, Capital Region of Denmark, Denmark; Center for Register Research, Institute of Economics, Aarhus University, Århus, Denmark The study was conducted under the auspices of the iPSYCH study (www.iPSYCH.au.dk)

**Keywords:** bi-genomic linkage disequilibrium, single nucleotide polymorphism, psychiatric disease

## Abstract

Mitochondria play a significant role in human diseases. However, disease associations with mitochondrial DNA (mtDNA) SNPs have proven difficult to replicate. A reanalysis of eight schizophrenia-associated mtDNA SNPs, in 23,743 normal Danes and 2,538 schizophrenia patients, revealed marked inter-allelic differences in haplogroup affiliation and nuclear ancestry, genogeophraphic affinity (GGA). This bi-genomic linkage disequilibrium (2GLD) could entail population stratification. Only two mitochondrial SNPs, m. 15043A and m. 15218G, were significantly associated with schizophrenia. However, these associations disappeared when corrected for haplogroup affiliation. The extensive 2GLD documented is a major concern when interpreting historic as well as designing future mtDNA association studies.

---

Genetic variants in mitochondrial DNA (mtDNA) – and in nuclear genes coding for mitochondrial function - have been associated with disease ^1-3^. More than 300 variants^4,5^ in mtDNA and genes involved in mitochondrial function^6^ have been reported to cause mitochondrial disease which is clinically characterised by complex metabolic, neurological, muscular and psychiatric symptoms^7,8^. SNPs in mtDNA and mitochondrial haplogroups (mtDNA hgs), which are evolutionarily fixed SNP sets with a characteristic geographical distribution, have been proposed as potential disease modifiers^8^. This has been reported in neurological degenerative diseases such as Alzheimer’s disease^9-12^ and Parkinson’s disease^12-14^, metabolic diseases and cancers^15^, as well as psychiatric diseases, notably schizophrenia (SZ) and bipolar disease^16-18^.

Association studies of mtDNA variants and disease have been difficult to replicate^8^. However, the definition of a methodological paradigm for association studies with mtDNA variants^19^ implicitly assumes that mtDNA variants are independent of the nuclear genome. In a recent Danish study on mtDNA haplogroups and their nuclear ancestry or nuclear genogeographical affinity (GGA), we demonstrated a marked difference in nuclear ancestry between individual haplogroups^20^. This means that mtDNA hgs entail population stratification also at the level of gDNA. The effect of such a stratification on disease association, will depend on the admixture structure of the particular population, the population history, epidemiology and genetic epidemiology of the disease, as well as the number of persons included in the study. The extensive fine-scale heterogeneity of gDNA and significant admixture documented in the UK^21^ and Europe^22^ further increase the risk of spurious false positive associations, if the mtDNA/gDNA interaction is not corrected phenomenon for in association studies.

Using DNA-array data from the Danish iPSYCH study on 2,538 schizophrenia patients and 23,743 population controls, we show that eight mtDNA SNPs, previously associated with SZ^16-18,23^, exhibit considerable inter-allelic differences both with respect to mtDNA hg affiliation and nuclear GGA. This phenomenon, that we name bi-genomic linkage disequilibrium (2GLD), affecting the association between an mtDNA SNP and mtDNA, as well as gDNA, can lead to both false negative and positive associations with disease. We demonstrate that for only two of the original eight SNPs is it possible to replicate the association with SZ in this cohort, when correcting for population stratification. Both mtDNA haplogroup affiliation and GGA affects the strength of association. Finally, we show that none of the SNPs are associated with SZ when examined on a particular the mtDNA hg background, with correction for 2GLD. As none of previous studies of mtDNA SNPs have been performed with correction for population stratification, let alone 2GDL, our results indicate that all such published associations should be considered preliminary. In principle, this conclusion should not be limited to associations with SZ.

## Results

From a literature search of mtDNA SNPs previously associated with SZ, we identified eight that were also typed by the PsychChip, Table 1. PsychChip data from 23,743 normal Danes and 2,538 SZ patients (Detailed in Suppl Table 1) showed that the SNPs were present in the population with frequencies varying from 0.2 % – 20.6 %, Table 1. There was no appreciable difference in mtDNA haplogroup or GGA distribution between controls and SZ patients, Suppl table 1.

**Table 1.** Schizophrenia associated mtDNA SNPs.

| Variant | Locus | Mol effect | Effect on risk of SZ | % DK iPSYCH controls (SZ) | Theoretical mtDNA hg associations* |
| --- | --- | --- | --- | --- | --- |
| m.1438A<sup>17</sup> | MT-RNR1 | n.a. | SZ↑ | 4.6 (4.8) | L0d, L1, C1b4, C4c2, M32'56, D5a2, I5c1, H1b1g, H2, H13a2b5, H14a2c, J1b1b1c, P2, P5 |
| m.3197C<sup>18</sup> | MT-RNR2 | n.a. | SZ↑ | 7.3 (8.2) | U5a'b, L3f1a1, H14b, U2e1a1a |
| m.3666A<sup>16</sup> | MT-ND1 | p.G120G | SZ↑ | 0.3 (0.4) | L0d1c1a1a, L0d1b2b2c2, L1, M13c, H1ak, R9b2, F1a1c, U5a2b3a1 |
| m.4769A<sup>17</sup> | MT-ND2 | p.M100M | SZ↑ | 4.4 (4.6) | R2, B4a1a1c, B4a1a4, L0d2b1, H2a |
| m.9377G<sup>16</sup> | MT-CO3 | p.W57W | SZ↓ | 0.5 (0.6) | G2a, Q2a1, D5b2, A2ac, H1c9a, U6a1b4, K1a4a1a2a, L2e, L3e2b1 |
| m.10398G<sup>23</sup> | MT-ND3 | p.T114A | SZ↑ | 20.6 (19.1) | K1,J, R11, B4m, B4c1c, R12'21, P4, U6a5c, K2a11, U8c, N1a1, W1e1a, W3a1d, N8, Y,S3,X2f1, R0a2k1 , B5 |
| m.15043A<sup>18</sup> | MT-CYB | p.G99G | SZ↑ | 5.7 (4.6) | M, N1a1, L2c2a, J1c2a3, T2f1a, U6a7, U2c1a |
| m.15218G<sup>18</sup> | MT-CYB | p.T158A | SZ↑ | 4.3 (5.2) | U5a1, M7a1a2, M10a1, HV1a'b'c, H13a2c |

\*Based on PhyloTree.

### Haplogroup distribution of mtDNA SNPs

The potential affiliation, based on PhyloTree, of SNPs to different mtDNA haplogroups is shown in Table 1, and the actual mtDNA haplogroup distribution in the controls (not different from that of the SZ patients, Suppl Table 2) is shown in Table 2. For all SNPs there is a marked difference in the actual mtDNA haplogroup distribution between the two alleles at the same position. Thus, when comparing persons with either of two alleles at the same mtDNA position, the comparison is between groups with widely differing mtDNA distributions. A PCA analysis of the mtDNA sequences in persons with either the A or G allele at position 15,043 is shown in figure 1A. This analysis shows that the difference in mtDNA sequence is very extensive.

**Table 2.**
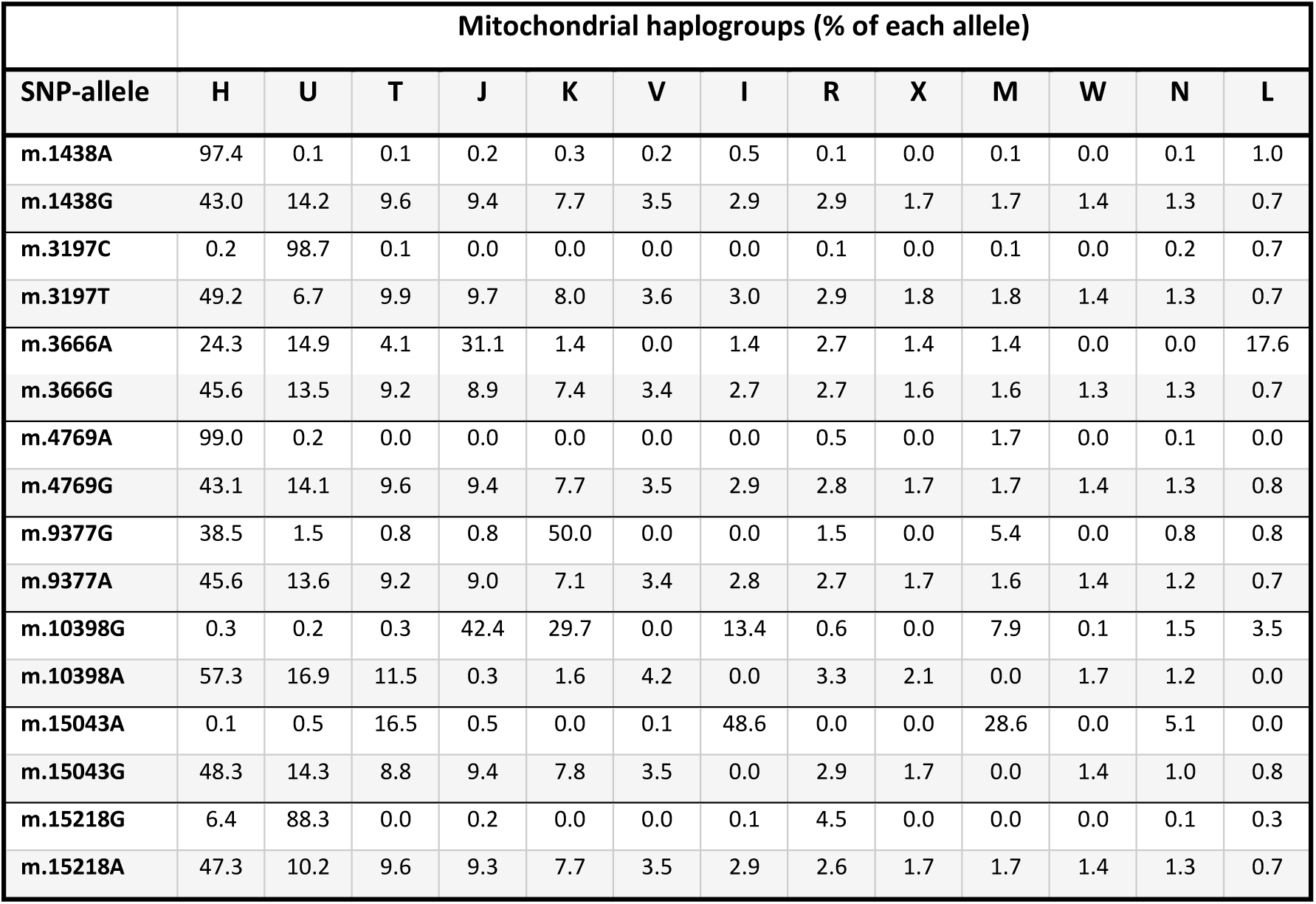
MtDNA haplogroup distribution for each SNP allele.

### Genogeographic affinity (ancestry) of mtDNA SNPs

The average GGA (ancestry) of the nuclear genome in control persons as a function of each mtDNA allele revealed major differences, both between different positions (inter-SNP) in the mtDNA molecule and between different alleles (inter-allelic) at the same position, Table 3. Apart from subtle differences in the distributions of Greenlandic and Asian GGA affiliations, the distribution is similar in SZ patients, Suppl table 3. Most marked is the difference in Danish GGA for m.15043 A/G, with an inter-allelic difference ~ 20 percent points in Danish ancestry, and the A-allele has a prominent Central South Asian ancestry not seen in the G-allele, Table 3 and Suppl Table 3. PCA analyses of the nuclear genome, Figure 1 B, of the two alleles, reveal a striking difference in the ancestry of the nuclear genome between the two alleles. Thus, when comparing persons with either of two alleles at the same mtDNA position, there is a substantial risk for comparison between groups with widely different ancestries (GGA), thus increased risk of confounding by population structure.

**Figure 1.**
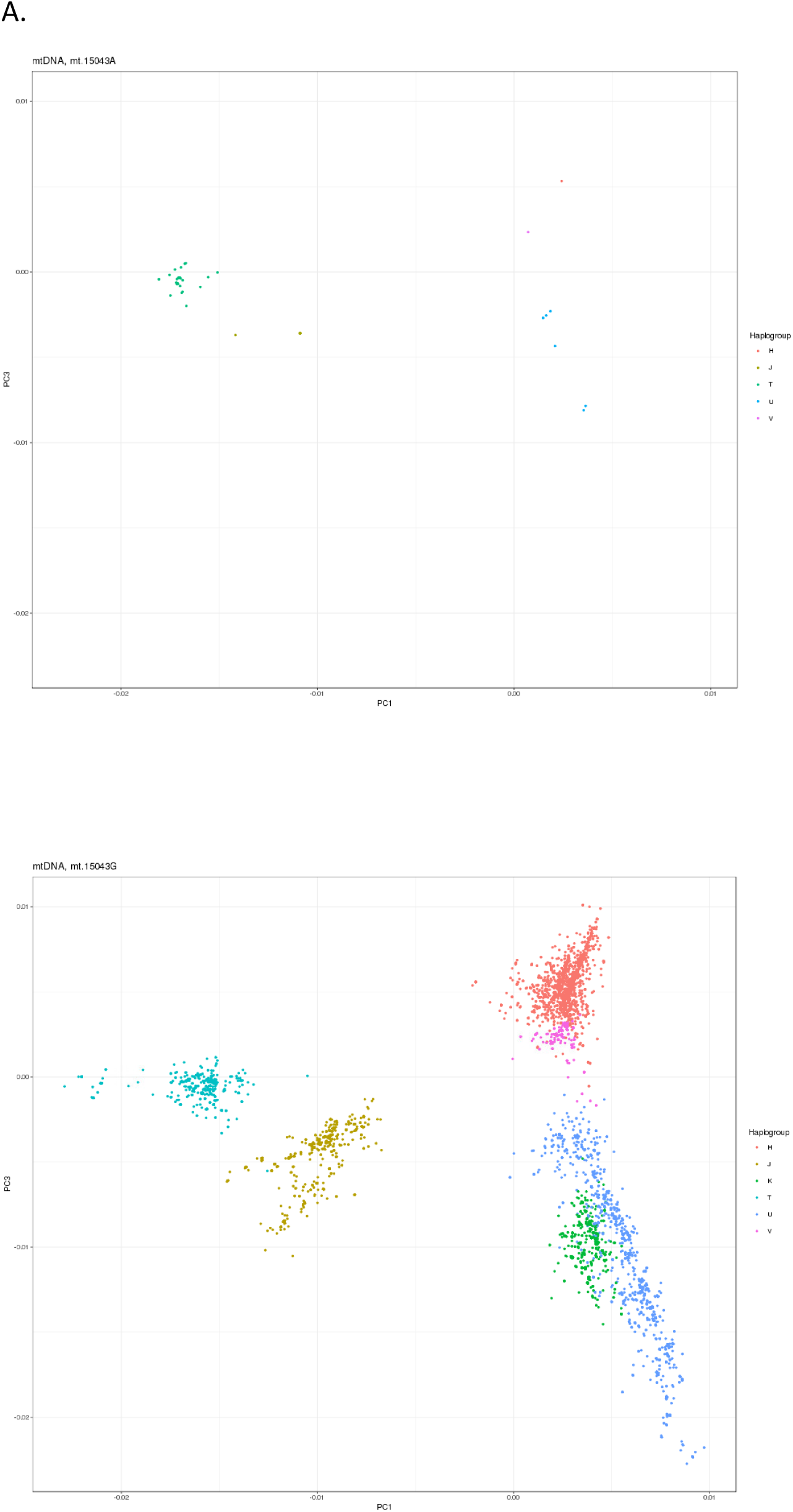
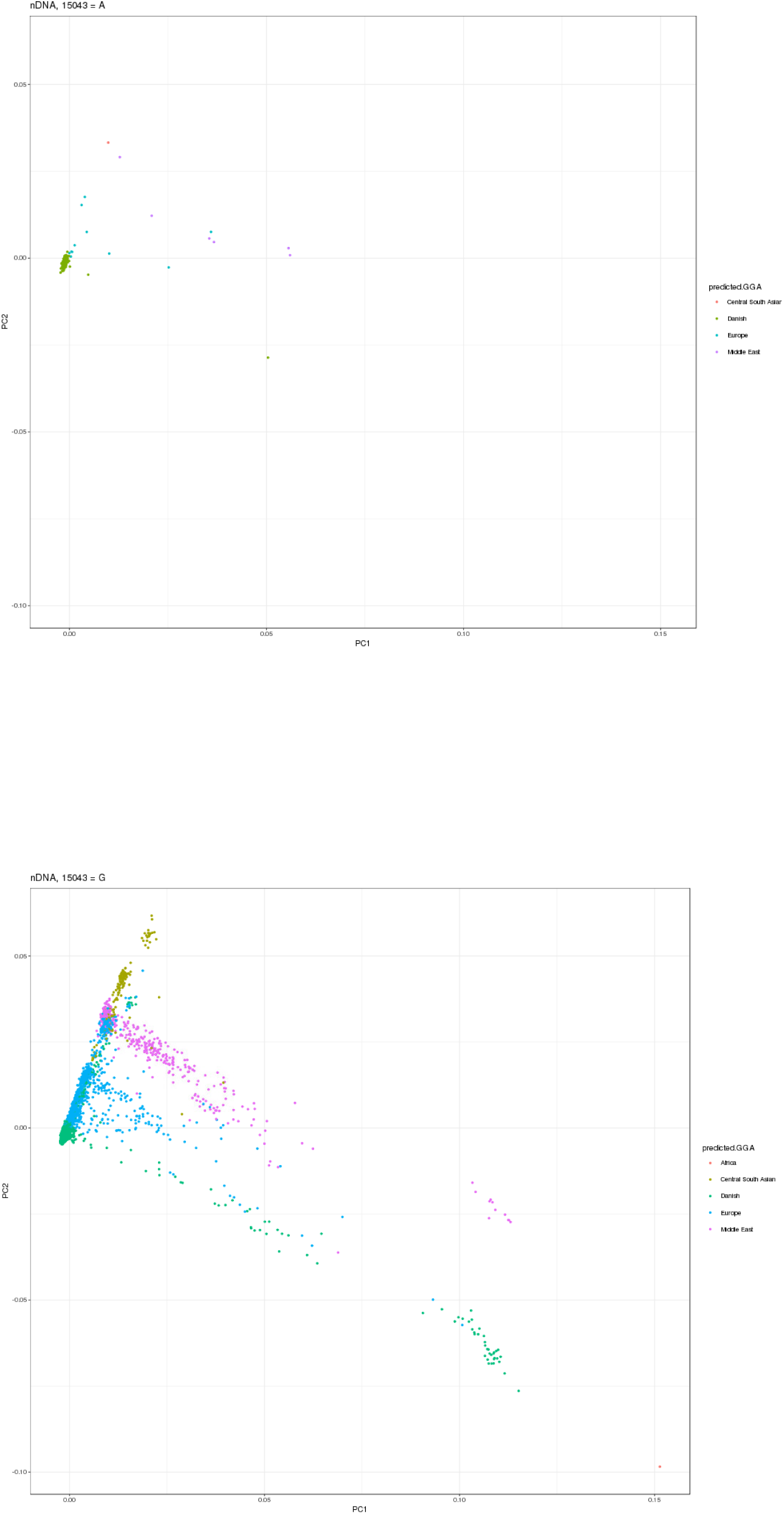
PCA of **A**. mtDNA (PC1 versus PC3) and **B**. nuclear genome (PC1 versus PC2), with the m.15043A (red) and m.15043G (blue) alleles. Only persons with “European” mtDNA haplogroup (H,HV,V,U,K,J & T) were included. The PCA was performed using PCs defined by control samples.

**Table 3.** Distribution of genogeographic affinities of controls with specific SNPs.

|  | Geographical origin (number of persons) |  |  |  |  |  |  |  |
| --- | --- | --- | --- | --- | --- | --- | --- | --- |
| SNP-allele | Denmark<br>(716) | Greenland<br>(592) | Europe<br>(161) | Middle<br>East<br>(178) | Central<br>South<br>Asia<br>(210) | East Asia<br>(241) | Africa<br>(101) | Oceania<br>(27) |
| m.1438A | 93.1 | 0.0 | 5.5 | 0.9 | 0.5 | 0.0 | 0.0 | 0.0 |
| m.1438G | 87.9 | 0.0 | 6.7 | 3.1 | 1.6 | 0.6 | 0.1 | 0.0 |
| m.3197C | 91.3 | 0.0 | 6.4 | 2.0 | 0.3 | 0.0 | 0.0 | 0.0 |
| m.3197T | 87.8 | 0.0 | 6.7 | 3.1 | 1.6 | 0.6 | 0.1 | 0.0 |
| m.3666A | 77.0 | 0.0 | 14.9 | 6.8 | 0.0 | 1.4 | 0.0 | 0.0 |
| m.3666G | 88.1 | 0.0 | 6.6 | 3.0 | 1.5 | 0.6 | 0.1 | 0.0 |
| m.4769A | 93.8 | 0.0 | 4.9 | 0.7 | 0.6 | 0.0 | 0.0 | 0.0 |
| m.4769G | 87.8 | 0.0 | 6.7 | 3.2 | 1.6 | 0.6 | 0.1 | 0.0 |
| m.9377G | 86.9 | 0.0 | 10.0 | 1.5 | 1.5 | 0.0 | 0.0 | 0.0 |
| m.9377A | 88.1 | 0.0 | 6.7 | 3.1 | 1.5 | 0.6 | 0.1 | 0.0 |
| m.10398G | 82.1 | 0.0 | 7.4 | 5.6 | 3.1 | 1.5 | 0.3 | 0.0 |
| m.10398A | 89.7 | 0.0 | 6.5 | 2.4 | 1.1 | 0.3 | 0.0 | 0.0 |
| m.15043A | 69.3 | 0.0 | 10.7 | 5.6 | 10.0 | 4.3 | 0.1 | 0.0 |
| m.15043G | 89.2 | 0.0 | 6.4 | 2.9 | 1.0 | 0.3 | 0.1 | 0.0 |
| m.15218G | 89.6 | 0.0 | 6.0 | 4.0 | 0.4 | 0.0 | 0.0 | 0.0 |
| m.15218A | 88.0 | 0.0 | 6.7 | 3.0 | 1.6 | 0.6 | 0.1 | 0.0 |

### Association between schizophrenia and mtDNA SNPs

The association of each mtDNA SNP with SZ was assessed (Table 4). In consequence of the inter-allelic differences in mtDNA hg affiliations and GGA demonstrated above, several association analyses were performed. The first (“ALL_hgs”) comprised all persons irrespective of either mtDNA hg or GGA, the second (“EU_hgs”) only comprised persons with a “classical European” mtDNA hg, i.e. H, V, J, K, U, or T. Thirdly, we examined associations in persons with a Danish GGA (All-hgs_DANES), and finally, in persons with both a “European” mtDNA haplogroup and a Danish GGA (EU-hgs_DANES). Five SNPs, m.1438A, m.3197C, m.3666A, m.4769A, and m.9388G showed no association with SZ, both when all persons were included and where selection was made to reduce effects of varying mtDNA and GGA affiliations. The m.10398G SNP was marginally significantly, while m.15043A was significantly associated with a reduced risk for SZ irrespective of the grouping. The high level of diversity in the PCA analysis of m.15043A (and the remaining SNPs – data not shown), figure 1A, prompted us to examine whether a fixation of the mtDNA hg, i.e. limiting the analysis to persons with a specific hg, in cases where it was reasonable frequent, would result in significant associations. The result, table 5, was the contrary – on a fixed mtDNA haplogroup background, none of the SNPs exhibited a significant association with SZ.

**Table 4.** Association of SNPs with SZ for different selections of cohort.

| Selection | mtDNA | Odds-ratio (95%cf. Interval) | P-Value |
| --- | --- | --- | --- |
| EU_hgs | 1438A | 1.04 (0.85-1.27) | 0.651 |
| ALL_hgs | 1438A | 1.04 (0.85-1.26) | 0.689 |
| EU_hgs_DANES | 1438A | 1.03 (0.83-1.26) | 0.755 |
| ALL_hgs_DANES | 1438A | 1.03 (0.84-1.27) | 0.716 |
| EU_hgs | 3197C | 1.13 (0.96-1.31) | 0.126 |
| ALL_hgs | 3197C | 1.13 (0.97-1.31) | 0.119 |
| EU_hgs_DANES | 3197C | 1.14 (0.97-1.34) | 0.101 |
| ALL_hgs_DANES | 3197C | 1.15 (0.98-1.35) | 0.085 |
| EU_hgs | 3666A | 1.33 (0.55-2.80) | 0.401 |
| ALL_hgs | 3666A | 1.27 (0.58-2.48) | 0.456 |
| EU_hgs_DANES | 3666A | 1.43 (0.59-3.05) | 0.377 |
| ALL_hgs_DANES | 3666A | 1.32 (0.54-2.79) | 0.402 |
| EU_hgs | 4769A | 1.04 (0.85-1.27) | 0.682 |
| ALL_hgs | 4769A | 1.06 (0.86-1.29) | 0.573 |
| EU_hgs_DANES | 4769A | 1.01 (0.82-1.23) | 0.874 |
| ALL_hgs_DANES | 4769A | 1.02 (0.82-1.26) | 0.832 |
| EU_hgs | 9377G | 0.89 (0.43-1.65) | 0.810 |
| ALL_hgs | 9377G | 1.05 (0.55-1.82) | 0.885 |
| EU_hgs_DANES | 9377G | 0.95 (0.46-1.77) | 1.000 |
| ALL_hgs_DANES | 9377G | 0.94 (0.46-1.75) | 1.000 |
| EU_hgs | 10398G | 0.97 (0.86-1.09) | 0.634 |
| ALL_hgs | 10398G | 0.91 (0.81-1.01) | 0.064 |
| EU_hgs_DANES | 10398G | 0.97 (0.86-1.10) | 0.686 |
| ALL_hgs_DANES | 10398G | 0.92 (0.82-1.03) | 0.153 |
| EU_hgs | 15043A | 0.94 (0.59-1.43) | 0.834 |
| ALL_hgs | 15043A | 0.80 (0.65-0.98) | 0.023 |
| EU_hgs_DANES | 15043A | 0.94 (0.58-1.46) | 0.913 |
| ALL_hgs_DANES | 15043A | 0.77 (0.60-0.97) | 0.029 |
| EU_hgs | 15218G | 1.27 (1.04-1.53) | 0.016 |
| ALL_hgs | 15218G | 1.25 (1.03-1.51) | 0.021 |
| EU_hgs_DANES | 15218G | 1.24 (1.01-1.52) | 0.037 |
| ALL_hgs_DANES | 15218G | 1.23 (1.01-1.50) | 0.044 |

**Table 5.** Association of SNPs with SZ in Danish GGA and a specific mtDNA haplogroup.

| **GGA** | **mtDNA hg** | **Allele** | **Odds-ratio (95% cfi)** | **p** | **SZ (N)** | **Controls (N)** |
| --- | --- | --- | --- | --- | --- | --- |
| Danish | H | m.1438A | 1.05 (0.84 – 1.29) | 0.67 | 110 | 1,000 |
| Danish | U | m.3197C | 1.08 (0.85 – 1.38) | 0.55 | 189 | 1,572 |
| Danish | H | m.3666A | 1.12 (0.13 – 4.71) | 0.70 | 2 | 17 |
| Danish | U | m.3666A | 2.59 (0.46 – 10.13) | 0.15 | 3 | 10 |
| Danish | J | m.3666A | 0.74 (0.08 – 3.06) | 1.00 | 2 | 22 |
| Danish | H | m.4769A | 1.03 (0.83 – 1.28) | 0.74 | 105 | 966 |
| Danish | H | m.9377A | 0.95 (0.38 – 3.07) | 0.81 | 1,046 | 9,917 |
| Danish | K | m.9377A | 0.90 (0.38 – 2.58) | 0.82 | 133 | 1,509 |
| Danish | H | m.10398A | 0.79 (0.18 – 7.14) | 0.67 | 1,049 | 9,947 |
| Danish | K | m.10398A | 1.40 (0.90 – 2.12) | 0.11 | 33 | 106 |
| Danish | T | m.15043A | 0.99 (0.59 – 1.58) | 1.00 | 22 | 208 |
| Danish | H | m.15218A | 1.09 (0.47 – 3.09) | 1.00 | 1,045 | 9,900 |
| Danish | U | m.15218A | 0.81 (0.63 – 1.04) | 0.09 | 211 | 1,939 |

## Discussion

Here we show that mtDNA SNPs are in bi-genomic linkage disequilibrium (2GLD) with mtDNA hgs and gDNA clusters due to population structure and shared demographic history of mtDNA and gDNA. This means that each mtDNA allele is associated with a unique distribution of mtDNA hgs and other associated mtDNA variants, and at the same time associated with a unique distribution of gDNA clusters. This has the consequence that an association between a particular allele in a specific SNP and disease is not exclusively a result of the presence of the particular allele of the SNP; rather it is the result of a combination of differences in mtDNA and gDNA sequences – a result of population stratification and admixtureIt is thus not – as is frequently done, and suggested as a paradigm for mtDNA association – sufficient to consider an association as evidence for a specific effect of an allele on the function of a protein or RNA coded for by the mtDNA and, consequently, as a cause of pathophysiological changes.

The linkage disequilibrium between different alleles on a SNP and mtDNA hgs and sub-hgs is not surprising as hgs are defined by series of evolutionarily conserved SNPs. The particular distribution of subsets of mtDNA hgs, sub-hgs and individual SNPs, that are associated with a particular allele at a specific SNP will depend on the population history and the extent and source of admixture. In most countries, and in particular in Europe, such history is very complicated and incompletely clarified. m.10398G was found associated with SZ in Han Chinese, however, when the cohort was broken down with respect to haplogroups, the association disappeared^24^. This illustrates that a specific allele’s mtDNA haplogroup distribution may induce spurious association with SZ. Spurious associations between mtDNA SNPs and a particular phenotype, when restricted to persons of a specific mtDNA hg^25^, may however be due to population stratification at the sub-hg level.

As mtDNA replicates without recombination, independently of the cell cycle, mtDNA hgs and SNPs should be independent of the nuclear genome, but only if the population were infinitely large and in the absence of population substructure. This is often not the case, due to geographical population substructure, recent admixture, socially and culturally defined restrictions in choice of spouse. A recent study showed that most Danish grandparents to present-day high school students chose spouses within a short distance of their birthplace^26^. This practice will with time lead to regionalisation, and a southwestern to northeastern gradient was found^26^. Furthermore, immigrants may seek a partner from within their ethnic community. Such effects have been sought eliminated in some studies by restricting the participants in mtDNA association studies to persons with a three-generation presence in the population. However, it has not been documented that this is sufficient to obviate association or linkage disequilibrium between mtDNA SNPs and specific gDNA clusters. Extensive gDNA micro-scale heterogeneity has been documented in the UK^21^ and Western France^27^ and admixture has been an important factor in the accretion of the present-day genomic variation of Europe^22,28^. The UK study^21^ showed that this is not just a result of recent demic changes; however, recent migrations may lead to widespread 2GLD.

Schizophrenia is a complex syndromic disease^29^ with geographically varying prevalence^30^ and characterized by a markedly elevated prevalence among first and second generation immigrants^31,32^, particularly among persons with dark skin moving to Nordic lattitudes^33^. These epidemiological characteristics of SZ obviously increase the risk of spurious associations caused by subtle admixture and 2GLD. However, it does not *per se* refute the mitochondrial pathogenic paradigm^34^ where variation in mitochondrial function, believed to interfere with ATP production 35,36, inflammation and signaling^37,38^ as well as Ca^2+^-homeostasis^39^, and apoptosis^38^, is considered to be of paramount importance for development of disease. Several neuroanatomical post-mortem findings in SZ brains indicate perturbed mitochondrial function^40^, but such findings are difficult to distinguish from changes caused by drug treatment.

The iPSYCH data are prospective and signs of immigration are apparent^20^, but they also showed that the variation in ancestry differed greatly between mtDNA hgs – even within traditional European hgs, i.e. mtDNA hg U, where ancient European sub-hgs occurred together with U-sub-hgs of recent Near Eastern and Central Asian origin^20^. Thus, 2GLD is likely to be a confounder and may lead both to false positive as well as false negative associations with disease. The method of correction for 2GLD in association studies will depend on the specific mtDNA SNP examined, the population structure and history, as well as the size of the study population.

If population stratification involving gDNA is inherent when performing association studies with mtDNA SNPs, it should be expected that diseases with geographically varying prevalence would be likely to find associated with specific mtDNA SNPs. The largest mtDNA association study to data^18^ found mtDNA SNPs associated with ulcerative colitis, exhibiting a European North-South and East-West gradient^41^, and with multiple sclerosis exhibiting a longitudinal prevalence gradient^42^ and effect of immigration^43^. The same study found that the prevalence of mtDNA SNPs associated with Parkinson’s disease was lower in African and Asian people^44^. Furthermore, the incidence of primary biliary cirrhosis is very high in North East England, 50 % lower in the rest of England and Scandinavia, and 90 % lower in the Middle East and Asia^45^.

A major problem with the interpretation of mtDNA SNP variants is the difficulties associated with performing a meaningful and reproducible assessment of mitochondrial function. In vitro studies of mitochondrial function, e.g. enzymatic activity measurements of OXPHOS components in cells, tissues^46^ or cybrids^47^ as well as allotopic expression^15^, are difficult to interpret as they also interfere with the inherent cellular control of mitochondrial function^38^. Furthermore, it should be noted, that mtDNA hgs and sub-hgs are cladistics groups and not functional units. Thus, in the Danish population, the U-hg is composed of a range of sub-hgs, e.g. U5a, U5b, U6, U7, and U8, with widely differing GGAs, reflecting migrations rather than selection^20^. It is thus meaningless to ascribe a specific functional effect to a particular mtDNA hg – without having carefully examined both mtDNA and nuclear genetic variation and corrected for stratifications in both.

Previous conflicting studies of disease associations with mtDNA have been suggested to be the result of insufficient power^48^, insufficient stratification respect to sex, age, geographical background^49^ or population admixture^50^, or the use of small areas of recruitment risking “occult” founder effects^51^. The fact that careful control, as here, of these factors and the 2GLD, results in none of eight previously SZ associated mtDNA SNPs being associated with SZ in the very large Danish iPSYCH cohort, suggests that previously reported associations could indeed be spurious findings due to cryptic population stratification. Meta-analyses pooling studies from different populations^15^ does not necessarily solve this problem – it may aggravate it by introducing further sub-stratification of the total population analysed. The extensive 2GLD demonstrated in the Danish population makes this phenomenon the most parsimonious explanation of non-replicable associations with mtDNA variants, not only for associations with SZ, but obviously 2GLD can interfere with associations between mtDNA and all types of diseases and traits.

## Online Methods

### Ethics statement

The iPSYCH cohort study (www.ipsych.au.dk) is register-based using data from Danish national health registries. The study was approved by the Scientific Ethics Committees of the Central Denmark Region (www.komite.rm.dk) (J.nr.: 1-10-72-287-12) and executed according to guidelines from the Danish Data Protection Agency (www.datatilsynet.dk) (J.nr.: 2012-41-0110). Passive, but not informed, consent was obtained, in accordance with Danish Law nr. 593 of June 14, 2011, para 10, on the scientific ethics administration of projects within health research.

### SZ patients and controls

As part of the iPSYCH recruitment protocol, 23,743 controls, born between May 1^st^ 1981 and Dec 31^st^ 2005 were selected at random from the Danish Central Person Registry. Among persons born within the same time span 2,538 persons assigned an ICD-10 F20 were identified in the Danish National Patient Registry. All were singletons, were alive one year after their birth, and had a mother registered in the Danish Central Person Registry. DNA should be available from DBS cards obtained from the Danish Neonatal Screening Biobank at Statens Serum Institut ^52^ Demographics of patients and controls are given in Suppl Table 1.

### Genetic analysis and mtDNA SNPing

From each DBS card two 3.2-mm disks were excised from which DNA extracted using Extract-N-Amp Blood PCR Kit (Sigma-Aldrich, St Louis, MO, USA)(extraction volume: 200 μL). The extracted DNA samples were whole genome amplified (WGA) in triplicate using the REPLIg kit (Qiagen, Hilden, Germany), then pooled into a single aliquot. Finally, WGA DNA concentrations were estimated using the Quant-IT Picogreen dsDNA kit (Invitrogen, Carlsbad, CA, USA). The amplified samples were genotyped at the Broad Institute (MA, USA) using the PsychChip (Illumina, CA, USA) typing 588,454 variants, developed by the Psychiatric Genetic Consortia. We then isolated the 418 mitochondrial loci and reviewed the genotype calls, before exporting into the PED/MAP format using GenomeStudio (Illumina, CA, USA). Haplo-grouping of mtDNA was performed using the defining SNPs reported in <u>www.phylotree.org</u> ^53^.

### Geno-geographic affinity (autosomal ancestry)

Geno-geographic affinity (GGA) estimation was done using ADMIXTURE 1.3.050 in the supervised approach. Briefly, reference populations consisting of Human Genome Diversity Project (HGDP) (http://www.hagsc.org/hgdp/), a Danish (716 individuals) and a Greenlandian (592 individuals) genotyping SNP data set were used. The final reference data set consisted of 103,268 autosomal SNPs and 2,248 individuals assigned to one of nine population groups: Africa, America, Central South Asia, Denmark, East Asia, non-Danish Europe, Greenland, Middle East and Oceania. The number of clusters, K was set to eight, based on principal component analysis clustering (data not shown). The subpopulations were merged with the reference population data set and analysed using ADMIXTURE. For prediction of the ancestry of individuals within the mtDNA haplogroups we created a random forest model^54^ based on the reference data set, with the clusters Q1-8 as predictors and population groups as outcome. Thus the ancestry analysis of the individual person was the result of a supervised prediction. Prediction was done using R3 version 3.2.2, using the Caret package.

### Statistics

The statistical significance of differences in mtDNA SNP proportions between controls and SZ patients was assessed using a permutation version of Fisher’s exact test. Samples with missing sequence data were excluded. Calculations were performed using R. Principal component analysis (PCA), was prepared using PLINK(v.1.90b3.31). For the PCA the reference population variants were extracted from the iPSYCH control sample, LD pruned (indep-pairwise 50 5 0.5) and allowing only SNPs with 99% genotyping rate. Prior to PCA of mtDNA data, samples were loaded into GenomeStudio (version 2011.a), a custom cluster was created using Gentrain (version 2), following automatic clustering, all positions with heterozygotes were manually curated. The data was exported relative to the forward strand using PLINK Input Report Plug-in (version 2.1.3). Eigenvectors were calculated using PLINK (v1.90b3.31). PCA plots were created using the package ggplot2 (version 1.0.1) in R (version 3.1.3).

## Acknowledgements

The iPSYCH study was funded by The Lundbeck Foundation Initiative for Integrative Psychiatric Research. We further gratefully acknowledge the financial support of The Jascha Foundation, The Strategic Research Council (“Heart Safe”), The Augustinus Foundation, The Lundbeck Foundation (Grant no. R67-A6552), and Familien Hede Nielsens Fond. This research has been conducted using the Danish National Biobank resource, supported by the Novo Nordisk Foundation.

## Legends to tables and figures

**Table 1.** The mtDNA alleles studied, their frequency, mtDNA haplogroup association, and link with disease.

**Table 2.** Distribution of mtDNA haplogroups of controls with specific mtDNA SNP.

**Table 3.** Autosomal genomic genogeographic affinity of studied mtDNA SNPs controls.

**Table 4.** Association between individual mtDNA SNPs and schizophrenia as a function of the selection of the involved persons. All hgs: All persons, EU hgs: persons with a European haplogroup; Danish GGA: persons with a Danish genogeographic affinity and Danish GGA and Eu hgs: persons both with a European hg and a Danish GGA.

**Table 5.** Association between mtDNA SNP and SZ in persons with Danish GGA and a defined mtDNA haplogroup.

